# The exotic species *Senecio inaequidens* pays the price for arriving late in temperate European grassland communities

**DOI:** 10.1101/542563

**Authors:** Benjamin M. Delory, Emanuela W.A. Weidlich, Miriam Kunz, Joshua Neitzel, Vicky M. Temperton

## Abstract

1. The South African ragwort (*Senecio inaequidens* DC.) is one of the fastest exotic plant invaders in Central Europe but, despite its large distribution area, it is still not commonly found in European grasslands. In order to better understand the mechanisms behind invasion resistance of grassland communities to *S. inaequidens*, we determined (1) to what extent the timing of arrival of *S. inaequidens* in the community affected its invasiveness as well as the performance of the native species, and (2) how the direction and strength of priority effects were affected by the composition of the native community being invaded, particularly with regard to the presence of N_2_-fixing species (legumes).
2. In a greenhouse experiment, we manipulated the timing of arrival of the exotic species in the community and the composition of the native background community to test the influence of these factors on the productivity and N content of exotic and native species. Using a set of interaction indices, we also investigated if the plant species origin (native or exotic) and the native community composition affected the benefit of arriving early and the cost of arriving late (i.e., priority effects) in the community.
3. We showed that both exotic and native species created inhibitory priority effects for late-arriving species. The establishment success of *S. inaequidens* strongly depended on its timing of arrival in a grassland community. On average, *S. inaequidens* benefited more from arriving early than the natives. We did not find any evidence to support that the presence of legumes in the background community would favour invasion by *S. inaequidens*. When natives arrived later than *S. inaequidens*, however, priority effects were weaker when legumes were present in the native community.
4. *Synthesis*: we showed that priority effects created by natives can lower the risk of invasion by *S. inaequidens* and are an important mechanism to explain why this exotic species is not commonly found in European grasslands yet. Our results suggest that an early arrival of this species at a site with low native species abundance (e.g., following a disturbance) is a scenario that could favour invasion by S. *inaequidens*.

## Introduction

Invasion of plant communities by non-native (“exotic” or “alien”) species is largely recognized as one of the main drivers of biodiversity loss worldwide (Sala *et al.* 2000; Elbakidze *et al.* 2018). In Europe, the number of alien vascular plant species has increased steadily since the beginning of the 19^th^ century, mainly because of enhanced economic activities such as trade and tourism increasing the risk of invasion (Elbakidze *et al.* 2018). Amongst all the exotic invasive plant species introduced to Europe, the South African ragwort (*Senecio inaequidens* DC., Asteraceae) is often considered as one of the fastest invaders (Lachmuth, Durka & Schurr 2010) and this is probably linked to its ability to colonize a wide range of ecological habitats (Heger & Böhmer 2005). *S. inaequidens* is a large perennial forb whose seeds have been repeatedly introduced to several locations in Central Europe via the transport of sheep wool imported from the Eastern highlands of South Africa (Ernst 1998; Lachmuth *et al.* 2010). It was observed for the first time in Europe (West Germany) at the end of the 19^th^ century (Ernst 1998; Heger & Böhmer 2005) and, after a time lag of nearly 80 years, it started to spread across Germany, mainly from a population introduced in Belgium at the beginning of the 20^th^ century (Lachmuth *et al.* 2010). It is an early-successional ruderal species requiring open sites with little resource competition, and is mainly found on disturbed sites and along railway tracks (Heger & Böhmer 2005). Despite its large distribution area in Central Europe and the fact that it has the potential to establish in native communities (Scherber, Crawley & Porembski 2003; Bossdorf, Lipowsky & Prati 2008), *S. inaequidens* is not commonly found in European grasslands and has not yet been considered as a threat to indigenous plant species (Heger & Böhmer 2005).

The invasion success of an exotic species in a new environment is multifactorial (Seastedt & Pyšek 2011). Most of the research in this area has been focusing on two complementary aspects of plant invasion. First, the identification of traits favouring the invasiveness of exotic species, such as high competitive ability (Sakai *et al.* 2001; Perkins & Hatfield 2014), germination and phenotypic plasticity (Allendorf & Lundquist 2003; Wainwright & Cleland 2013), the ability to reproduce vegetatively/clonal growth (Kolar & Lodge 2001), or the ability to produce allelochemicals that supress local species at new sites (Callaway & Aschehoug 2000; Callaway & Ridenour 2004; Aschehoug *et al.* 2014). Second, the identification of native plant community characteristics affecting their susceptibility to invasion (invasibility), such as the disturbance regime (Chytrý *et al.* 2008; Seastedt & Pyšek 2011), fluctuations in resource availability (Davis, Grime & Thompson 2000; Liu, Zhang & van Kleunen 2018), species and functional group richness (Tilman 1997; Knops *et al.* 1999; Naeem *et al.* 2000; Wardle 2001; Kennedy *et al.* 2002; Fargione & Tilman 2005; Pokorny *et al.* 2005; Scherber *et al.* 2010; Mason, French & Jolley 2017) species and functional group composition (Crawley *et al.* 1999; Prieur-Richard *et al.* 2002; Fargione, Brown & Tilman 2004; Wardle *et al.* 2011; Byun, de Blois & Brisson 2013; Yannelli *et al.* 2017), and the presence of natural enemies (Keane & Crawley 2002; Shea & Chesson 2002; Levine, Adler & Yelenik 2004). Because exotic species often germinate more quickly, grow faster, and take up resources more efficiently than native species (Wainwright, Wolkovich & Cleland 2012; Wilsey, Barber & Martin 2015), the invasion process is also tightly linked to the concept of priority effect in ecology, in which the species arriving first at a site significantly affect the development, growth, and reproduction of species arriving later (Chase 2003; Vannette & Fukami 2014; Temperton *et al.* 2016).

In grasslands, priority effects caused by biotic interactions can have effects that supersede abiotic influence on the community. Priority effects caused by species arriving before others can affect community structure as well as ecosystem functioning both aboveground (Wilsey *et al.* 2015; Weidlich *et al.* 2017) and belowground (Körner *et al.* 2008; Weidlich *et al.* 2018). Such priority effects occur either because the early-arriving species reduce the amount of resources available for late-arriving species (called niche preemption) (Fukami 2015), or because the early-arriving species modified the type of niches available for the species arriving later via, for instance, extra nitrogen (N) availability if N_2_-fixing species arrive first, root exudation or the selection of a particular soil microbiome (called niche modification, including plant-soil feedbacks) (Callaway *et al.* 2004; Suding *et al.* 2013; van der Putten *et al.* 2013; Perkins & Hatfield 2014; Fukami 2015). Depending on the timing of arrival of an exotic species and the desired outcome (resistance of the community to invasion), priority effects can be considered as positive or negative. During ecological restoration of degraded landscapes, for example, there is much potential for creating positive priority effects by sowing natives before the arrival of exotics (Hess, Mesléard & Buisson 2019). If exotic species with strong competitive abilities are given a head-start (e.g., after a disturbance), however, they can quickly outcompete native species (Abraham, Corbin & D’Antonio 2009; Grman & Suding 2010; Stevens & Fehmi 2011; Dickson, Hopwood & Wilsey 2012; Wainwright *et al.* 2012; Ulrich & Perkins 2014; Wilsey *et al.* 2015; Stuble & Souza 2016), and such priority effects can persist for several years (Martin & Wilsey 2012; Vaughn & Young 2015).

To what extent the timing of arrival of *S. inaequidens* affects its capacity to invade a European grassland community is not yet known but, considering that (1) it is an early-colonizing species able to produce a large amount of wind-dispersed seeds (Ernst 1998), (2) it has the capacity to establish and survive in mature grassland communities (Scherber *et al.* 2003), and (3) introduced *S. inaequidens* populations respond to greater resource availability by increasing aboveground and belowground productivity and reproductive output (Bossdorf *et al.* 2008), it is likely that an early arrival of this species (i.e., before natives) could lead to successful invasion of grassland communities.

Both direct and indirect facilitation between invading exotic species (Simberloff & Von Holle 1999; Flory & Bauer 2014) or between natives and exotics (Bruno, Stachowicz & Bertness 2003; Bulleri, Bruno & Benedetti-Cecchi 2008; Saccone *et al.* 2010) are important invasion mechanisms. When looking at the effect of natives on exotics, facilitation in the form of N fertilization by leguminous species as well as nurse plant interactions have been shown to increase communities’ susceptibility to invasion (Maron & Connors 1996; Mason, French & Jolley 2013). With regard to grassland ecosystems, the presence of legumes is known to increase the amount of N available for its neighbours via two co-occurring mechanisms: (1) direct N transfer (N transfer) from legumes to non-legume neighbours, and (2) reduced interspecific competition for soil mineral N (N sparing) when legumes derive most of their N from the atmosphere (Temperton *et al.* 2007). How such facilitation mechanisms affect the invasiveness of an exotic species has been the central issue in many research studies, with some showing that the presence of N_2_-fixing species in the community can favour invasion (Prieur-Richard *et al.* 2000, 2002; Mwangi *et al.* 2007; Scherber *et al.* 2010), and others not showing such positive relationship between invasibility and legume presence (Tilman 1997). In contrast, the presence of strong competitors, such as grasses, tends to increase the resistance of a plant community to invasion (Prieur-Richard *et al.* 2000, 2002; Fargione *et al.* 2004; Mwangi *et al.* 2007; Scherber *et al.* 2010). As described above, both the species and functional group composition of a native plant community can have important consequences for invasion but, surprisingly, little is known about the effect of this factor on the establishment of *S. inaequidens* in European grasslands. Because N facilitation can be expected when legumes are present in a community, we hypothesize that grassland communities containing N_2_-fixing species would be more susceptible to invasion by *S. inaequidens* than communities containing only grasses. In other words, we expect to find weaker priority effects of native species on *S. inaequidens* if legumes are present in the community being invaded (i.e., *S. inaequidens* benefits from the presence of legumes that arrived earlier than it did). Similarly, when the exotic species is the first to arrive in the community, we also expect to find weaker priority effects acting on the late-arriving native species if legumes are present (i.e., the non-legumes in the community will benefit from N facilitation and thus suffer less asymmetric competition).

In this paper, we present the results of a controlled greenhouse experiment designed to evaluate the benefits of arriving early and the costs of arriving late (i.e., priority effects) for the exotic species *S. inaequidens* invading a community composed of species characteristic of mesotrophic temperate European grasslands. The costs and benefits for native species arriving earlier or later than *Senecio* were also evaluated in this study. In addition to time of arrival, we also manipulated the composition of the native background community to test if this would affect the strength and direction of priority effects for both native and exotic species. Here, we addressed the following questions:

1. Does the invasiveness of *S. inaequidens* in a European grassland community depend on its timing of arrival and the composition of the native background community?
2. Is the performance of the native species affected by the timing of arrival of *S. inaequidens* and the composition of the native community?
3. Does *S. inaequidens* benefit more than the native species from arriving early in the community?
4. Does the composition of the native plant community affect the direction and strength of priority effects for exotic and native species?
5. Do the native species pay a greater cost than *S. inaequidens* when arriving late in the community?

## Material and Methods

### Plant material

All seeds used for the experiment described in this paper were native wild species, not cultivars, provided by Rieger-Hofmann GmbH (Blaufelden-Raboldshausen, Germany).

### Experimental design

We set up an experiment using a full factorial and randomized design to test the influence of two main factors on the performance of an exotic (*S. inaequidens*) and native species characteristic of temperate mesotrophic European grasslands: (1) the timing of arrival of the exotic species in the plant community, and (2) the composition of the native plant community. With regard to the timing of arrival of the exotic species, it was sown either earlier, later, or at the same time (synchronous) as the native species. Because manipulating the timing of arrival of the exotic species implied the organisation of two sowing events (sowing interval: 21 days), two synchronous treatments were set up so that all priority effect treatments could be directly compared to plants that had grown for the same length of time: one at the first sowing (Synchronous 1), and another one at the second sowing (Synchronous 2). Therefore, the factor timing of arrival of the exotic species comprised a total of four levels (Fig. 1). Two different native plant communities were used in this experiment. In the first scenario, *S. inaequidens* was sown into a community composed of grasses only (*Holcus lanatus, Festuca pratensis, Phleum pratense*). In the second scenario, it was sown into a community composed of grasses (*H. lanatus, F. pratensis, P. pratense*) and legumes (*Medicago sativa, Trifolium pratense, Lotus corniculatus*). Each of the 8 treatment combinations (4 timing of arrival of *S. inaequidens* × 2 native plant community compositions) was replicated 5 times. This experiment was set up in a greenhouse located in Rotes Feld, Lüneburg (Lower Saxony, Germany). During the period of the experiment, the temperature inside the greenhouse was 23.5 ± 5.2 °C during the day and 16.8 ± 2.8 °C during the night.

**Figure 1.**
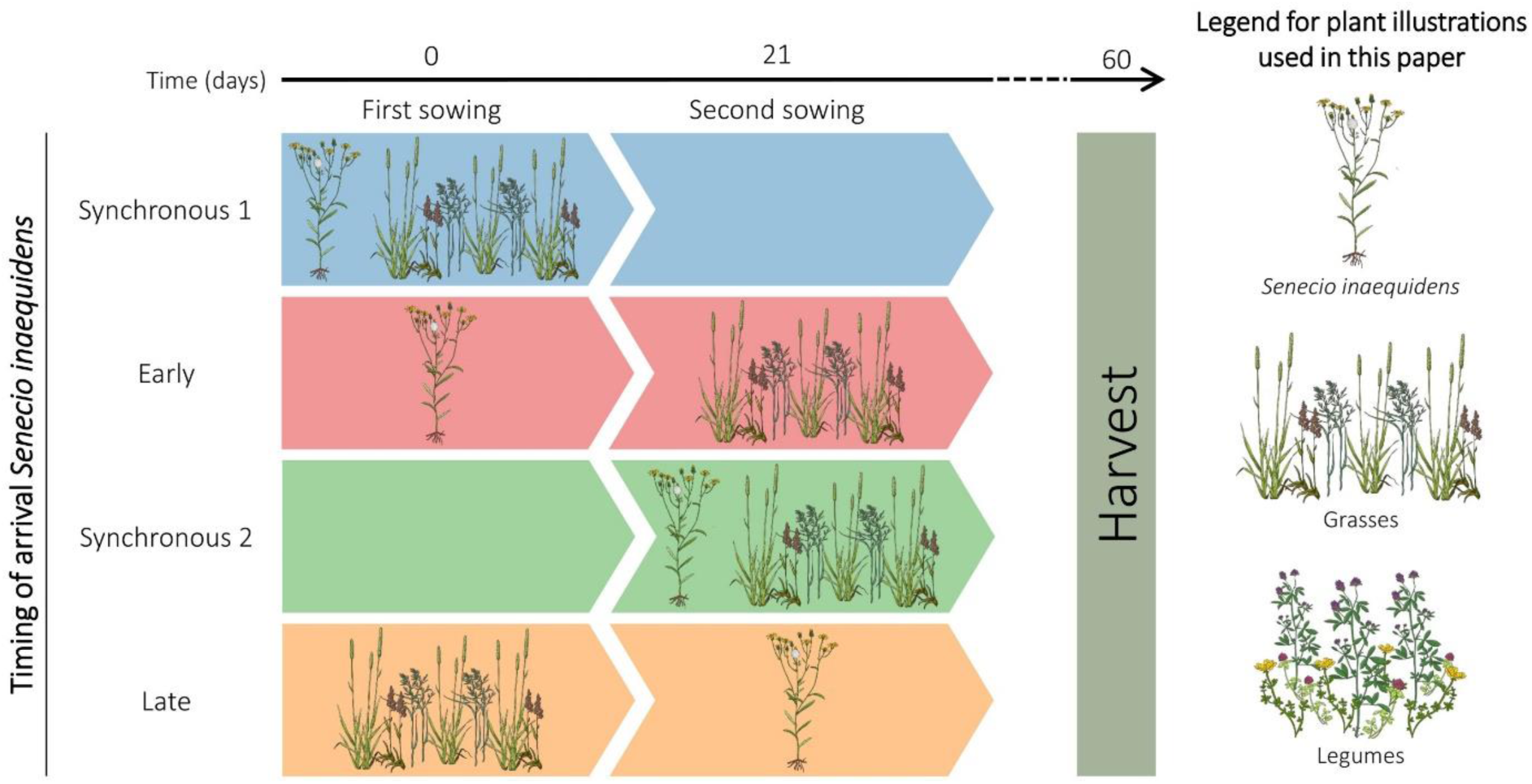
Manipulating the timing of arrival of the exotic species *S. inaequidens* in a community composed of native species. Note that, as shown in this figure and all subsequent figures, the terms “late” or “early” always refer to the timing of arrival of the exotic species, not the background community. For clarity reasons, this figure only illustrates one of two possible scenarios with regard to the composition of the native plant community invaded by *S. inaequidens* (only grasses). A second scenario in which *S. inaequidens* was sown a community composed of grasses and legumes was also tested. A legend for plant illustrations used throughout this paper is provided on the right side of the figure. Plant illustrations by Carolina Levicek.

Eight days before the start of the experiment, 5 L of sand was added at the bottom of 40 pots (volume: 18.4 L, top surface area: 784 cm^2^, height: 30 cm). Then, the pots were filled with a mixture of sand (50%, v/v) and potting soil (50%, v/v) up to the top. After filling, the pots were placed inside a greenhouse and were watered twice before the start of the experiment. We started the experiment by sowing the species arriving first in the communities (Fig. 1). Depending on the timing of arrival of the exotic species, we sowed either *S. inaequidens* (Early arrival), the native species (Late arrival), or both *S. inaequidens* and the native species (Synchronous 1). Twenty-one days later, we sowed the rest of the species in the pots (Fig. 1). During this second sowing event, we sowed either the native species (Early arrival), *S. inaequidens* (Late arrival), or both *S. inaequidens* and the native species (Synchronous 2). After each sowing event, a thin layer of sand and potting soil (50%/50%, v/v) was added at the top of the pots to favour germination. For each treatment, the sowing density of each species was adjusted to take into account differences in plant composition and germination rates so to allow for an even community outcome. When the native community was composed of grasses only, the sowing density was adjusted to reach a target of 50 individuals/species growing in the pots. When the native community was composed of a mixture of grasses and legumes, however, the sowing density was adjusted to reach a target of 25 individuals/species. For all treatments, the sowing density of the exotic species was calculated to reach a target of 25 individuals growing in each pot. The composition and sowing densities used for each plant community are summarized in Table S1. All pots were regularly watered using tap water throughout the duration of the experiment.

Sixty days after the first sowing event, aboveground biomass was harvested separately for *S. inaequidens*, the legumes, and the grasses. Shoot samples were then dried at 70 °C until constant mass was reached.

### Measurements

For each pot, we measured the total shoot dry weight of *S. inaequidens*, legumes, and grasses. The total carbon (%C) and nitrogen (%N) content in *S. inaequidens,* legume, and grass shoots (the latter were pooled into grass or legume biomass) was determined with a C/N analyser (Vario EL; Elementar, Langenselbold, Germany) using 14.7 ± 1.5 mg of dry and finely ground material. Evidence for N facilitation in communities containing legumes was investigated using the δ^15^N natural abundance method (*sensu* Temperton *et al.* 2007). Sample δ^15^N values (‰) were calculated using equation 1, where R represents the ratio of ^15^N/^14^N isotopes. *R*_*sample*_ values were determined using an elemental analyser (Elementar Vario EL Cube) coupled to a stable isotope ratio mass spectrometer (IR-MS, Isoprime). Isotope ratios were determined for *S. inaequidens*, grasses, and legumes using dried and finely ground plant material (*S. inaequidens*: 7.6 ± 1.1 mg; grasses: 6.7 ± 0.1 mg; legumes: 9.0 ± 0.1 mg). Atmospheric N_2_ is the international standard used for IR-MS measurements of δ^15^N (*R*_*standard*_). If legumes were actively fixing atmospheric N_2_, *R*_*sample*_ values measured in their shoots should be very close to the *R*_*standard*_ value measured in the atmosphere. Therefore, the expected δ^15^N value measured in the shoots of actively fixing legumes should be close to zero. For species relying solely on soil N, however, we expect δ^15^N values to be different from zero (in theory, close to the soil δ^15^N value) because soil N often has a greater ^15^N abundance than atmospheric N_2_ (Unkovich *et al.* 2008). Apparent N transfer between legumes and non-legume neighbours was assessed by comparing the δ^15^N values measured in *S. inaequidens* and grass shoots growing with or without legumes in the community. If belowground N transfer occurred, we expect non-legume shoots to have lower δ^15^N values when growing in communities containing leguminous species (Temperton *et al.* 2007). Because non N_2_-fixing species can also benefit from the soil N that is not taken up by leguminous species (N sparing), we used both the N status (%N) and the δ^15^N values measured in plant shoots to investigate how N dynamics was affected by the timing of arrival of the exotic species and the species composition of the native community.

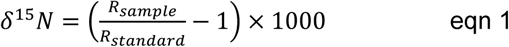

### Quantification of priority effects

In assembly research, the strength of priority effects has mainly been quantified using interaction indices in the form of log response ratios (Vannette & Fukami 2014; Stuble & Souza 2016). Because such indices are not bounded between finite values, they are however not well suited for comparing results between different experiments (Díaz-Sierra *et al.* 2017). Here, we introduce a set of standardized, symmetric, and bounded interaction indices designed to quantify the benefit of arriving early (*B*) and the cost of arriving late (*P*) during community assembly (Table 1; note that in our definition of priority effects, these only occur as effects on later arriving species). These indices share the same mathematical properties as the relative interaction index commonly used to measure competition and facilitation between interacting plants, i.e. they are standardized, symmetric around zero, and are bounded between −1 and +1 (Díaz-Sierra *et al.* 2017). The direction of the priority effect is given by the sign of *P*, with inhibitory priority effects having negative values, and facilitative priority effects having positive values. The strength of the priority effect is given by the absolute value of *P*. Because the calculation of *B* and *P* relies on the comparison of the performance of organisms arriving at different time in the community, but having the same age at harvest (e.g. *Senecio* growing in Early and Synchronous 1 treatments; see Fig. 1), they can only be calculated if the experiment includes as many synchronous treatments (i.e., all species arriving at the same time) as sowing events. The values for *P* and *B* reported in this paper were all calculated using shoot dry weight data. The total biomass of native species in the background community was calculated by summing the biomass of legumes and grasses. In the supplementary material, cost and benefit values calculated using shoot N content data are also provided (Fig. S4).

**Table 1.** Quantification of priority effects. In the equations, *Y* is a particular response variable (e.g., biomass production, N content, etc.). The superscripts and subscripts refer to the timing of arrival of the exotic species and the origin of the plant species on which *Y* was measured (*E* is for exotic, *N* is for natives), respectively.

|  | For the **exotic**<br>species | For the **native**<br>species | Interpretation |  |  |
| --- | --- | --- | --- | --- | --- |
|  |  |  | Negative<br>values | 0 | Positive values |
| **Benefit (B) of**<br>**arriving early** | $B_E = \frac{Y_E^{Early} - Y_E^{Sync1}}{Y_E^{Early} + Y_E^{Sync1}}$ | $B_N = \frac{Y_N^{Late} - Y_N^{Sync1}}{Y_N^{Late} + Y_N^{Sync1}}$ | Negative effect<br>of early arrival | No<br>benefit | Positive effect<br>of early arrival |
| **Cost (P) of**<br>**arriving late** | $P_E = \frac{Y_E^{Late} - Y_E^{Sync2}}{Y_E^{Late} + Y_E^{Sync2}}$ | $P_N = \frac{Y_N^{Early} - Y_N^{Sync2}}{Y_N^{Early} + Y_N^{Sync2}}$ | Inhibitory<br>priority effect | No<br>priority<br>effect | Facilitative<br>priority effect |

### Statistical analyses

Generalised linear models (GLMs) were used to investigate the effect of the timing of arrival of the exotic species (Arrival), the species composition of the native plant community (Composition), and their interaction (Interaction) on the aboveground biomass productivity of the exotic species and the grasses. Because the number of *S. inaequidens* individuals that established varied from pots to pots, we tested if adding the number of *Senecio* individuals that were actually growing in the pots as a covariate would improve the quality of the models. After comparing models fitted with and without the number of *Senecio* individuals, we did not find any evidence to support that models accounting for the number of *Senecio* individuals were better than the models that did not account for it. In addition, there was no correlation between the number of *Senecio* individuals growing in a pot and the total productivity achieved by the exotic species (r=0.153, P=0.346). Therefore, in this paper, we only report the results of statistical models fitted without using the number of *Senecio* individuals growing in the pots as a covariate. When the interaction term did not significantly improve the model, a new model without interaction term was fitted. GLMs were also used to investigate the effect of the timing of arrival of the exotic species on the aboveground productivity of the legumes. GLMs were always fitted on plant biomass data using a Gamma distribution and a log-link function.

The effect of the timing of arrival of the exotic species, the species composition of the native plant community, and their interaction on the C/N content and δ^15^N values measured in *S. inaequidens* and grass shoots was investigated using two-way ANOVA models. One-way ANOVA models were used to test for the effect of the timing of arrival of the exotic species on the C/N content and δ^15^N values measured in legume shoots.

Two-way ANOVA models were used to test if the plant species origin (native or exotic), the native community composition, and their interaction had an effect on the benefit of arriving early (*B*_*E*_ or *B*_*N*_) and the cost of arriving late (*P*_*E*_ or *P*_*N*_) in the community. When reported, 95% confidence intervals were computed by bootstrapping (1000 iterations) using the percentile method. We considered that the mean value of a group was not significantly different from zero when the 95% confidence interval of that group included zero.

General linear hypotheses (post-hoc tests) were tested using Tukey contrasts and the glht function of the multcomp R package (Hothorn, Bretz & Westfall 2008). When results of post-hoc tests are shown, adjusted P-values (single-step method) are reported to account for multiple comparisons of group means. All statistical analyses were performed in R 3.4.3 (R Core Team 2018) with an alpha value of 0.05.

## Results

### Timing of arrival and native plant community composition affect the performance of *S. inaequidens*

When *S. inaequidens* was the first species to arrive in a community, it produced significantly more biomass than when arriving at the same time as the native species (Fig. 2A). The effect of the timing of arrival of *Senecio* on plant productivity, however, varied with the species composition of the native community (Interaction: F_1,16_=5.31, P=0.035). If the native plant community contained both grasses and legumes, the biomass gain due to an early arrival of *Senecio* (+137.7%) was larger than when the native community contained only grasses (+64.1%). This result can be explained by the fact that, when both exotic and native species arrived at the same time, *S. inaequidens* was less productive in communities containing legumes than in communities containing only grasses (Fig. 2A; z=2.96, P=0.012), while the productivity achieved by *Senecio* did not differ between both communities when it arrived earlier than the native species (Fig. 2A; z=-0.15, P=0.998). Whatever the composition of the native plant community, an early arrival of *S. inaequidens* did not affect its N content (Fig. 2C; Arrival: F_1,16_=1.19, P=0.291; Composition: F_1,16_=0.37, P=0.553; Interaction: F_1,16_=4.33, P=0.054). With regard to the C content of *S. inaequidens* shoots, it was negatively affected by an early arrival of *Senecio* (−3.8%; Arrival: F_1,16_=9.29, P=0.008; Composition: F_1,16_=1.13, P=0.304; Interaction: F_1,16_=1.88, P=0.190), but only when the native community was composed of legumes and grasses (Fig. S1-A; t=3.12, P=0.023).

**Figure 2.**
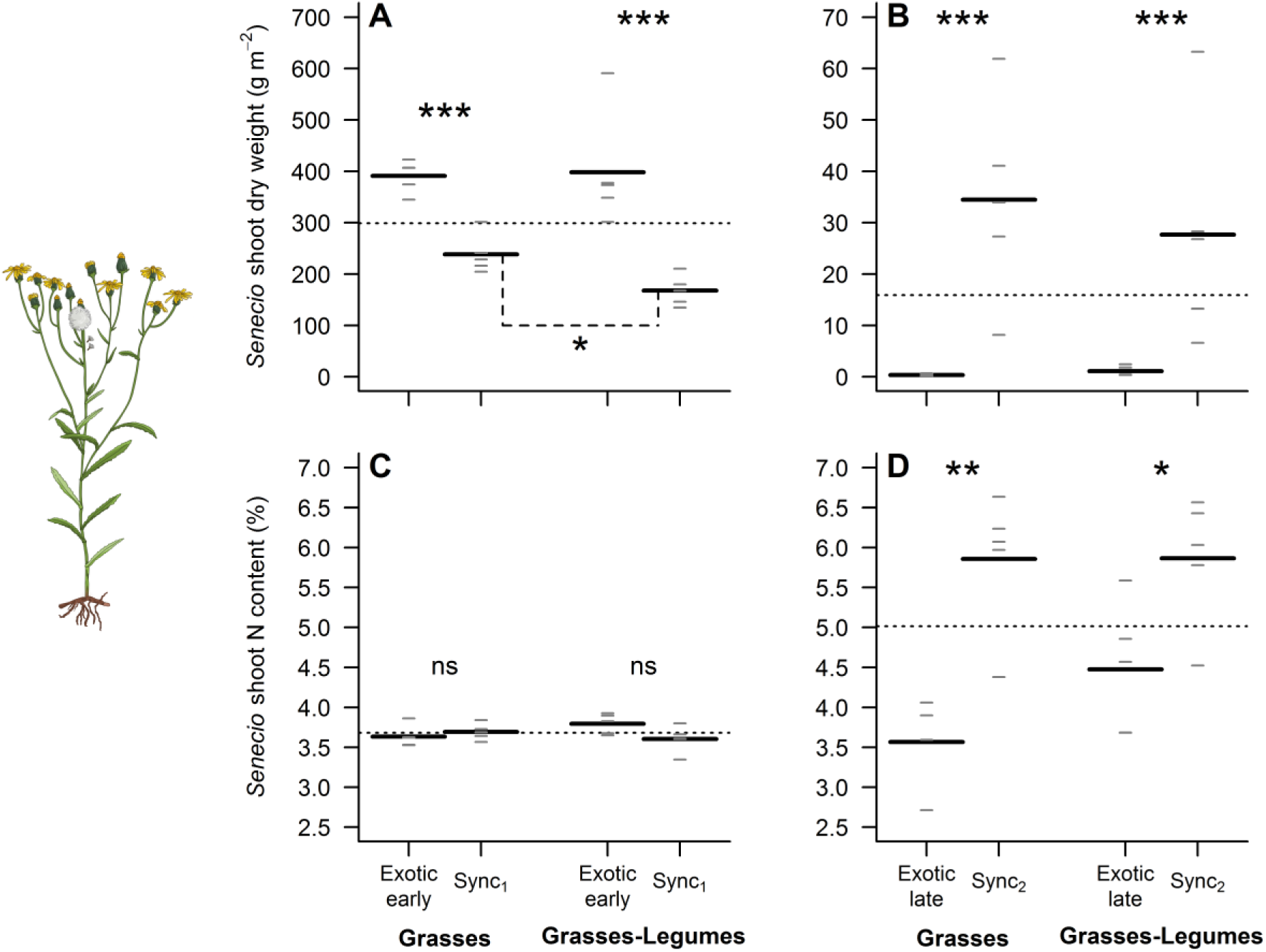
Timing of arrival and native community composition affects the performance of *S. inaequidens*. All panels show the overall mean (horizontal dotted line), the mean of each group (long black horizontal lines, n=4-5), and each individual observation (short grey horizontal lines). Note that the terms “late” or “early” in all graphs always refer to the timing of arrival of the exotic species (Fig. 1). Because of the low shoot dry weight of *S. inaequidens* obtained for one replicate of the treatment where the exotic species arrived late in a community made of native grass species only, we were not able to measure the N content of that replicate, leaving n=4 in this case. ns, not significant (P>0.05); *, P<0.05; **, P<0.01; ***, P<0.001.

When the exotic species arrived later than the natives, it barely managed to establish and produced between 96.1% and 99.0% less biomass than the plants that arrived at the same time as the native species (Fig. 2B; Arrival: F_1,16_=62.74, P<0.001). When it was the last species to arrive, the productivity achieved by *Senecio* did not depend on the composition of the native community (Composition: F_1,16_=1.16, P=0.297; Interaction: F_1,16_=3.19, P=0.093). For both plant community compositions, a late arrival of *S. inaequidens* had a strong negative effect on its shoot N content (Fig. 2D; Arrival: F_1,15_=24.90, P<0.001). Although we did not find any significant effect of the species composition of the native plant community on the N content of *S. inaequidens* (Fig. 2D; Composition: F_1,15_=0.83, P=0.377; Interaction: F_1,15_=1.53, P=0.234), we observed that the decrease in N content associated with a late arrival of the exotic species was lower in communities containing legumes (−23.7%) than in communities containing only grasses (−39.1%). For both native community compositions, the C content of *S. inaequidens* shoots was similar whether it arrived later or at the same time as the native species and did not differ between the two plant communities (see supplementary Fig. S1-B; Arrival: F_1,15_=10^-4^, P=0.993; Composition: F_1,15_=0.94, P=0.348; Interaction: F_1,15_=1.74, P=0.207).

### Timing of arrival of the exotic species and plant community composition affect the biomass production and N content of native grass species

In comparison with a situation where both native and exotic species arrived at the same time, the aboveground productivity of the grass species was between 62.2% and 66.0% greater if they arrived earlier than the exotic (Fig. 3A; Arrival: F_1,16_=41.61, P<0.001), and this result was independent of plant community composition (Interaction: F_1,16_=0.02, P=0.885). When the natives arrived before the exotic species, the timing of arrival of the exotic species was the only factor affecting the shoot N content (Fig. 3C; Arrival: F_1,16_=5.12, P=0.038; Composition: F_1,16_=1.27, P=0.276; Interaction: F_1,16_=0.48, P=0.497). On average, the grasses had a lower N content when they arrived earlier than the exotic species. This difference in N content was not significant anymore when the effect of the timing of arrival of the exotic species was investigated separately for each plant community (Fig. 3C).

**Figure 3.**
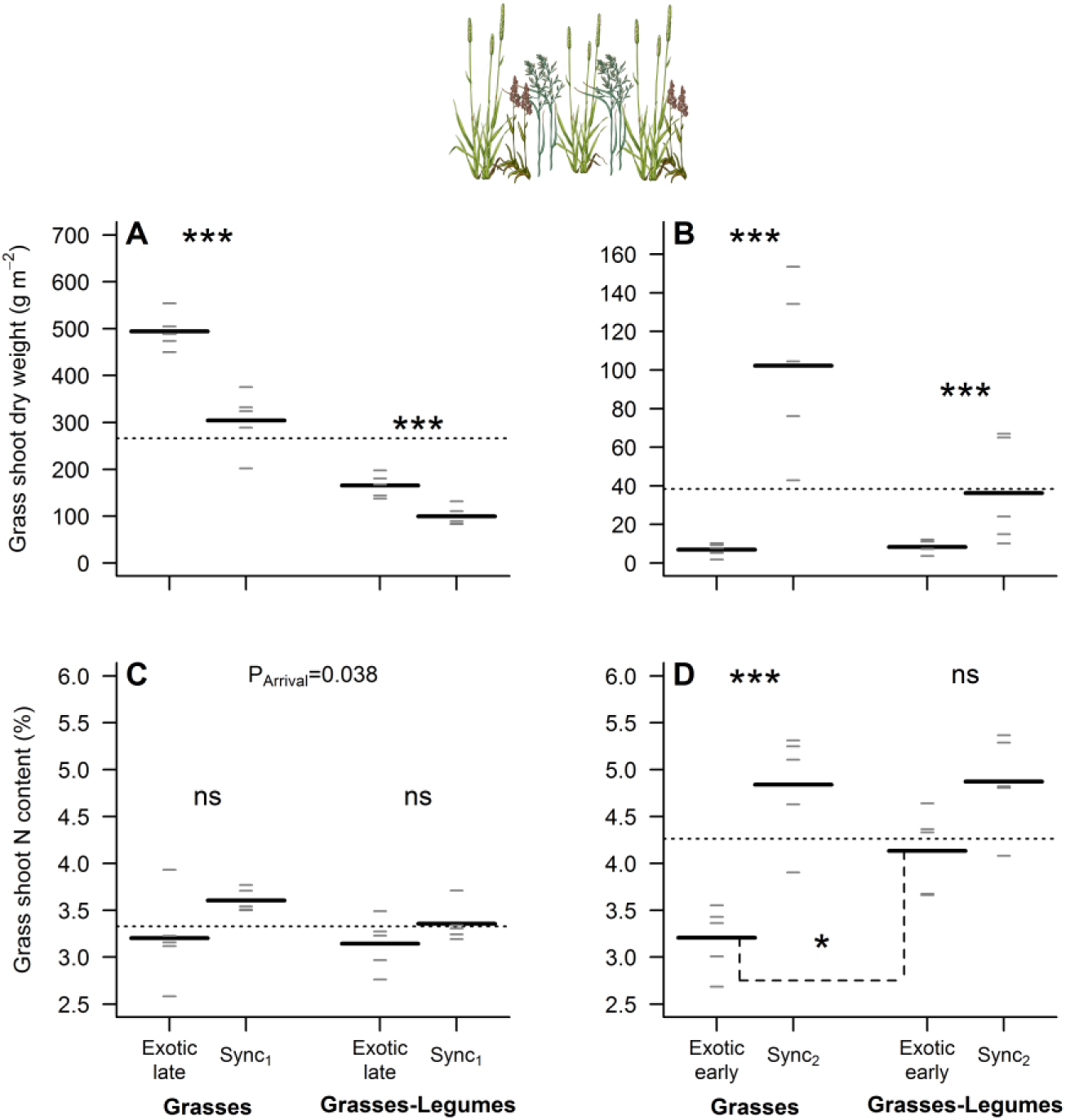
Biomass production and N content of native grass species are affected by the timing of arrival of the exotic species and the composition of the native plant community. All panels show the overall mean (horizontal dotted line), the mean of each group (long black horizontal lines, n=5), and each individual observation (short grey horizontal lines). Note that the terms “late” or “early” in all graphs always refer to the timing of arrival of the exotic species, such that the “late” treatment outcome shows the performance of grasses when they arrive early (Fig. 1). ns, not significant (P>0.05); *, P<0.05; **, P<0.01; ***, P<0.001.

Overall, the grasses had a hard time to establish when they arrived later than the exotic species. In this situation, they produced between 77.2% and 93.3% less biomass than the grasses that arrived at the same time as *S. inaequidens* (Fig. 3B). This timing of arrival effect, however, was dependent on the native community composition (Interaction: F_1,16_=4.84, P=0.043), mainly because the lower grass density in communities containing legumes led to a lower grass biomass production in these communities when all species arrived at the same time (Fig. 3B). On average, the grasses arriving later than *S. inaequidens* also had a significantly lower N content (−24.4%) than the grasses that arrived at the same time as the exotic species, particularly when native communities did not contain any legumes (Fig. 3D; Arrival: F_1,16_=30.18, P<0.001; Composition: F_1,16_=4.96, P=0.041; Interaction: F_1,16_=4.31, P=0.054). For both native community compositions, the N content of grass shoots was lower when the native species arrived after the exotic, but the difference with the synchronous treatment was only significant when legumes were absent (Fig. 3D; t=5.35, P<0.001). In addition, when *Senecio* was the first species to arrive, the N content in grass shoots was 28.9% greater if the native community included legumes (Fig. 3D; t=-3.04, P=0.026).

Neither timing of arrival of the exotic species, nor the species composition of the native community affected the C content of grass tissues (Fig. S2).

### Timing of arrival of the exotic species affects the biomass production and N content of native legume species

When legumes arrived before *S. inaequidens*, they were 43.6% more productive than when arriving at the same time as the exotic species (Fig. 4A; F_1,8_=27.87, P<0.001). We did not observe any effect of a late arrival of the exotic species on the N content (Fig. 4C; F_1,8_=0.03, P=0.867) and C content (Fig. S3-A; F_1,8_=0.88, P=0.376) of legume shoots.

**Figure 4.**
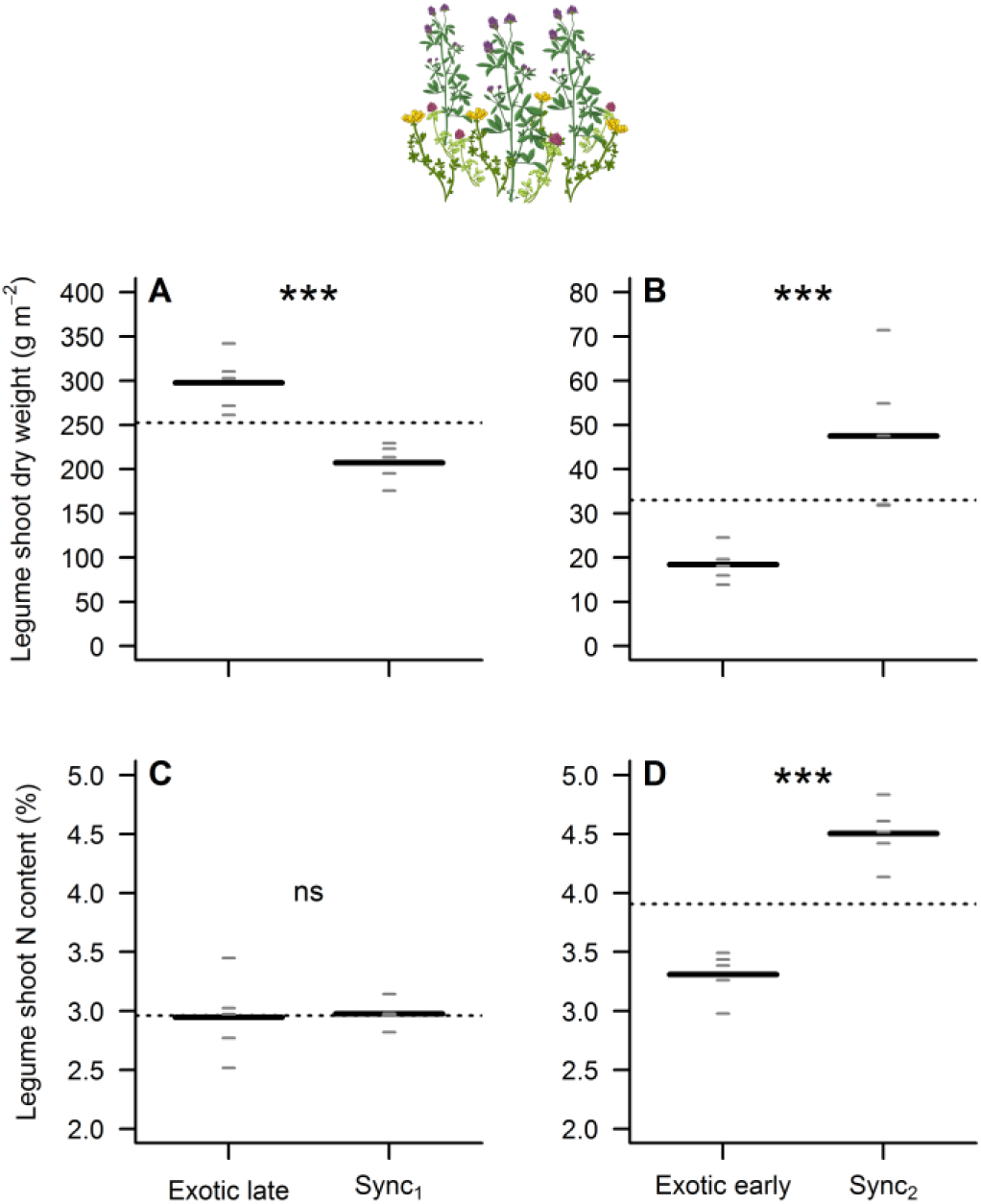
Biomass production and N content of native legume species are affected by the timing of arrival of the exotic species. All panels show the overall mean (horizontal dotted line), the mean of each group (long black horizontal lines, n=5), and each individual observation (short grey horizontal lines). Note that the terms “late” or “early” in all graphs always refer to the timing of arrival of the exotic species, such that the “late” treatment outcome shows the performance of legumes when they arrive early (Fig. 1). ns, not significant (P>0.05); *, P<0.05; **, P<0.01; ***, P<0.001.

When *S. inaequidens* was the first species to arrive in the community, the aboveground productivity and the shoot N content of the legumes decreased by 61.2% (Fig. 4B; F_1,8_=25.54, P<0.001) and 26.5% (Fig. 4D; F_1,8_=66.36, P<0.001) in comparison with control plants sown at the same time as the exotic species, respectively. An early arrival of the exotic species did not have any effect on the C content (Fig. S3-B; F_1,8_=0.01, P=0.913) of legume tissues.

### *S. inaequidens* benefits more from arriving early than the natives

Our results showed that both exotic and native species benefited from arriving early in the community but, on average, *S. inaequidens* benefited more than the natives (Fig. 5A; Species: F_1,16_=5.56, P=0.031). Interestingly, the effect of the native community composition on the benefit of arriving early differed between exotic and native species (Fig. 5A; Interaction: F_1,16_=5.31, P=0.035). The positive effect associated with an early arrival was greater for the exotic species if it was followed by a mixture of grasses and legumes compared with a mixture of grass species only (t=2.56, P=0.040). For the natives, however, the benefit of arriving early was similar for both community compositions (t=-0.70, P=0.741). Very similar results were obtained when the benefit of arriving early was calculated using shoot N content data (Fig. S4).

**Figure 5.**
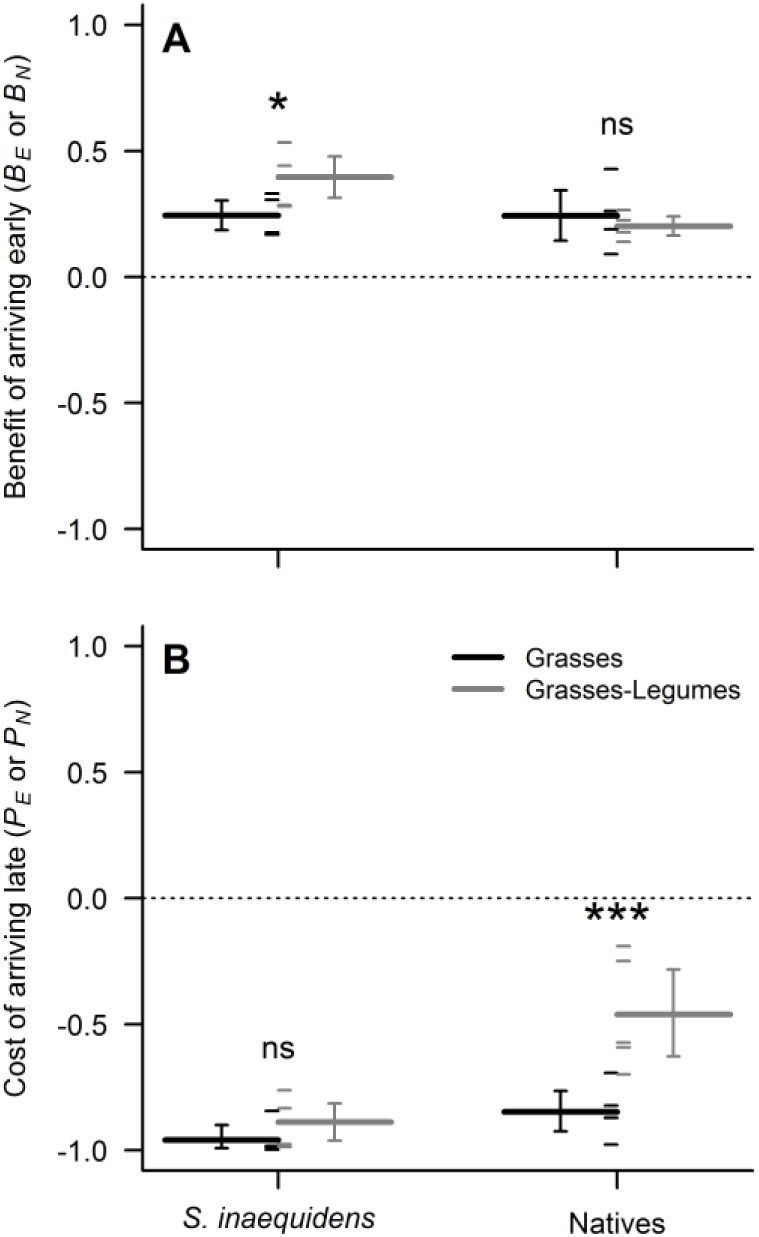
Benefit of arriving early and cost of arriving late in the community for exotic and native species. Benefits and costs were calculated based on shoot dry weight data using the equations listed in Table 1. All panels show the mean of each group (long horizontal lines, n=5), and each individual observation (short horizontal lines). Results are shown separately for each plant species origin (exotic or natives) and each native community composition (see legend). Error bars are 95% confidence intervals computed by bootstrapping using the percentile method. If a 95% confidence interval does not include zero, the mean value of the group is significantly different from zero. ns, not significant (P>0.05); *, P<0.05; **, P<0.01; ***, P<0.001.

### Arriving late is less costly for natives than *S. inaequidens*, particularly when legumes are present in the community

Both exotic and native species created inhibitory priority effects for species arriving later (Fig. 5B). Overall, growth reduction by priority effects was strongest for the exotic species (Species: F_1,16_=19.45, P<0.001) and when legumes were absent from the plant community (Community: F_1,16_=13.97, P=0.002). In addition, the composition of the native community affected differently the priority effect strength measured on exotic and native species arriving later (Interaction: F_1,16_=6.63, P=0.020). Our results showed that the strength of priority effects acting on *S. inaequidens* did not depend on the composition of the native community it tried to invade (t=0.82, P=0.660). The strength of priority effects acting on native species, however, was significantly affected by the composition of the native community. In comparison with a scenario where the native community is composed of grasses only, inhibitory priority effects acting on natives following an early establishment of the exotic species were on average 45.7% weaker when legumes were present in the native community (t=4.46, P<0.001). Very similar results were obtained when the strength of priority effects was calculated using shoot N content data (Fig. S4).

### Evidence for atmospheric N_2_ fixation by legumes, but not for N transfer to non-legume neighbours

On average, the δ^15^N values measured in legume shoots were 47.6% and 42.2% lower than those measured in *S. inaequidens* and grass shoots, respectively (Fig. 6A). This result strongly suggests that the legumes harvested at the end of the experiment were actively fixing atmospheric N_2_. However, the positive δ^15^N values measured in legume shoots also indicate that the legumes did not rely solely on the fixation of atmospheric N_2_ as a source of N for plant growth, but also on the soil N pool.

**Figure 6.**
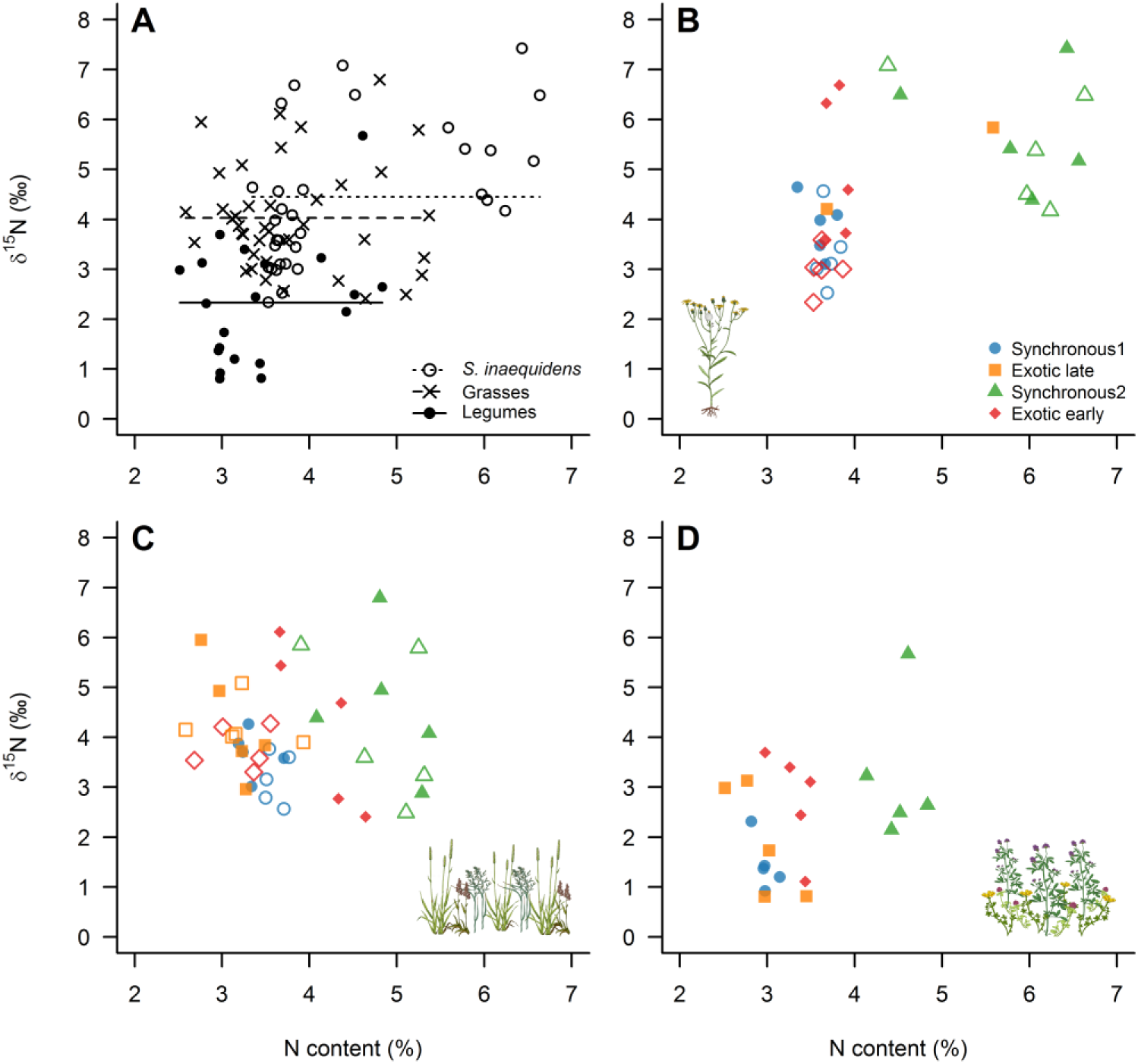
Relationship between the N content and the δ^15^N values measured in *S. inaequidens* (A, B), grass (A, C), and legume shoots (A, D). Panel A shows all data together. The horizontal lines represent the mean δ^15^N values calculated for *S. inaequidens*, the grasses, and the legumes (see legend in panel A). In panels B-D, dots are coloured based on the timing of arrival of the exotic species (see legend in panel B). Open symbols refer to native communities containing only grasses, while closed symbols refer to native communities containing grasses and legumes. Because of the low shoot dry weight of *S. inaequidens* obtained for one replicate of the treatment where the exotic species arrived late in a community made of native grass species only, we were not able to measure the N content of that replicate, giving n=4 for this treatment. For the same reason, we were not able to measure the δ^15^N values in all replicates of the treatment where *S. inaequidens* arrived late in a community composed of grass species only, as well as in three replicates of the treatment where *S. inaequidens* arrived late in a community composed of grasses and legumes.

The δ^15^N values measured in the exotic species were significantly affected by its timing of arrival (Arrival: F_2,24_=11.54, P<0.001) and by the composition of the native community being invaded (Composition: F_1,24_=6.19, P=0.020), but not by the interaction between these two factors (Interaction: F_2,24_=2.11, P=0.143) (Fig. 6B). Surprisingly, *S. inaequidens* shoots had greater δ^15^N values when legumes were present in the community (t=3.10, P=0.005). In addition, *S. inaequidens* shoots from the Synchronous 2 treatment also had greater δ^15^N values than the ones harvested from the Early (t=3.94, P=0.002) and Synchronous 1 treatments (t=3.41, P=0.006).

When looking at the native species, the δ^15^N values measured in grass shoots were neither affected by the timing of arrival of the exotic species (Arrival: F_3,32_=1.59, P=0.212), nor by the species composition of the native plant community (Composition: F_1,32_=1.17, P=0.287; Interaction: F_3,32_=0.11, P=0.954) (Fig. 6C). Similarly, we did not find any evidence to support that the timing of arrival of the exotic species affected the δ^15^N values of legume shoots (Arrival: F_3,16_=2.86, P=0.070) (Fig. 6D).

Despite that legumes were actively fixing atmospheric N_2_ during our experiment, the fact that the shoot δ^15^N values measured in *S. inaequidens* and native grass species were not lower when grown in communities containing legumes did not support direct N transfer from legume to non-legume neighbours.

## Discussion

The results presented here confirmed that the timing of arrival of *S. inaequidens* has a strong impact on its performance. This exotic species performed much better (greater biomass production) when it was the first species to arrive in the community. In fact, pots in which the exotic species was sown before the natives were near monocultures, with *S. inaequidens* accounting for 96% of the total aboveground productivity, while it accounted for only 40% of the total biomass when it was sown at the same time as the native species. On the other hand, it barely managed to establish (less than 0.6% of the total aboveground productivity) and had a lower N content when it arrived later. The timing of arrival of the exotic species in the community also strongly affected the performance of the native species. When the natives arrived three weeks before *Senecio*, they produced more biomass than natives arriving at the same time as the exotic species. When the natives arrived later in the community, however, they had a hard time to establish (lower biomass production) and had a lower shoot N content. Therefore, whatever the origin of the plant species arriving first (either exotic or native), it created strong inhibitory priority effects for the species arriving later in the assembly process. These results are in line with previous studies that have shown that either positive (lower abundance of exotic species) or negative (natives outcompeted by exotics) priority effects can be created depending on whether the exotic species arrive later or earlier than the natives, respectively (Abraham *et al.* 2009; Grman & Suding 2010; Stevens & Fehmi 2011; Dickson *et al.* 2012; Wainwright *et al.* 2012; Ulrich & Perkins 2014; Wilsey *et al.* 2015; Stuble & Souza 2016).

Using nectar-inhabiting microorganisms as a model system, Vannette & Fukami (2014) demonstrated that priority effects are stronger if (1) species arriving at different time in the community use resources in a similar way (niche overlap), (2) early-arriving species have a strong impact on the local environment (impact niche), and (3) the growth, survival, and reproduction of late-arriving species is greatly affected by environmental conditions (requirement niche). Because both exotic and native species created strong inhibitory priority effects in our experiment, it is likely that the species that were sown first impacted the environment in such a way that it degraded the requirement component of the late-arriving species’ niches. In an attempt to classify the mechanisms behind the creation of priority effects during community assembly, Fukami (2015) distinguished two main categories of mechanisms by which early arriving species impact their local environment and affect the establishment of species arriving later. The first mechanism, referred to as niche preemption, is mainly resource-driven. It is based on the assumption that the plant species arriving first at a site affect species arriving later by reducing the amount of available resources such as light, water, and soil nutrients. The second mechanism, referred to as niche modification, is based on the assumption that the species arriving first in the community will modify the types of niches available locally and will affect the identity of the species that will be able to colonize the community. Contrary to niche preemption mechanisms, which can only lead to the creation of inhibitory priority effects, niche modification mechanisms can lead to the creation of both facilitative (e.g., soil fertilization by leguminous species) or inhibitory priority effects (e.g., exudation of allelochemicals altering the growth rate of species arriving later) (Maron & Connors 1996; Callaway & Aschehoug 2000; Fukami 2015). Which class of mechanisms led to the creation of priority effects in our experiment is not clear yet, but the near competitive exclusion and lower N content of species arriving late in the community strongly suggest that niche preemption mechanisms played an important role (Fukami 2015). In addition, we did not find any evidence of apparent N transfer from legumes to non-legume neighbours in our experiment, thus not supporting niche modification by leguminous species via N fertilization.

*S. inaequidens* being an early-successional species (Ernst 1998; Heger & Böhmer 2005), it is likely to be one of the first species to arrive at open sites, particularly in disturbed and stony areas. Our results showed that *S. inaequidens* benefited more from arriving early in the community than the native species, and this effect was particularly strong when it was followed by a mixture of grasses and legumes. This result is in agreement with the hypothesis that, due to their earlier emergence, greater germination rates, and faster growth, exotics would benefit more than native species from arriving early in the community (Dickson *et al.* 2012; Wainwright *et al.* 2012; Wilsey *et al.* 2015; Hess *et al.* 2019). This result, however, contradicts other priority effect studies that showed that exotic species benefited equally (Stuble & Souza 2016) or less (Cleland, Esch & Mckinney 2015) than natives when they were the first to arrive in a community. As suggested by Stuble & Souza (2016), differences between studies could arise from differences between species and study systems (e.g., testing annuals vs perennials).

Overall, *S. inaequidens* suffered more from arriving late than the native species. This key result contradicts previous studies that showed that the cost of arriving late in a plant community tends to be lower for exotics than for natives (Stuble & Souza 2016), or that exotics create stronger priority effects than natives (Wilsey *et al.* 2015). Arriving late was less costly for the native species than for the exotic species, suggesting a possible evolutionary adaptation of native grasslands species to finding free niches despite high canopy cover of the community into which they are trying to establish. Interestingly, our results also showed that the strength of the priority effects acting on the exotic species did not depend on the composition of the native community being invaded. Contrary to our expectations, the growth inhibition of *S. inaequidens* when it arrived in a community composed of native grasses only was as strong as when it arrived in a mixture of grasses and legumes. Because (1) the native grass community had a greater N content per unit plant biomass than the community composed of a mixture of grasses and legumes (Fig. S5), and (2) the two native communities did not differ in productivity across our priority effect treatments (Fig. S6), the grass community seemed better at taking up soil N than the grass-legume mixture in our experiment.

Even though the amount of soil N available for plant growth was probably greater in the community containing legumes, the fact that the strength of the priority effects acting on *S. inaequidens* was not different between the two native communities used in our experiment strongly suggests that available soil N was not the limiting factor for the establishment of the exotic species in our artificial grassland communities. Instead, preemption of light or other soil resources by natives might be more important mechanisms to explain the near competitive exclusion of late-arriving *S. inaequidens* invaders (Ernst 1998; Heger & Böhmer 2005; Frankow-Lindberg 2012; Wilsey *et al.* 2015). The composition of the native plant community, however, had a strong impact on the priority effects created by the exotic species on late-arriving natives. When a mixture of native grasses and legumes followed *S. inaequidens*, these priority effects were nearly 50% weaker than when legumes were absent from the native community. Although we found evidence for N sparing in communities containing legumes, as shown by the greater N content of late-arriving *S. inaequidens* or grass species when legumes were present (Fig. 6), we cannot fully conclude that the decrease in priority effect strength observed for the grass-legume mixture was solely due to N facilitation associated with the presence of N_2_-fixing species in the community, mainly because the two native community compositions used in this study differed in species and functional group richness.

There is now an expanding body of literature claiming that the creation of priority effects would be a useful technique to restore degraded habitats, alter competitive relationships, and steer plant communities towards desirable states in terms of biodiversity and functioning (Wilsey *et al.* 2015; Temperton *et al.* 2016; Weidlich *et al.* 2017, 2018; Young *et al.* 2017). Manipulating plant community assembly to promote native species that will ultimately exert strong priority effects on exotics is also a very interesting approach to lower the risk of invasion (Hess *et al.* 2019). Most of the research in this field has been performed in the USA (Young *et al.* 2017; Goodale & Wilsey 2018). The main reason for this is that American grasslands suffer more from invasion by exotic species than European ones (Seastedt & Pyšek 2011). To the best of our knowledge, this study is one of the few that explicitly tested how historical contingency by priority effects impact on the establishment of a rapidly expanding exotic species in European grasslands (Lang *et al.* 2017). Priority effects being contingent on environmental conditions during plant establishment (Young *et al.* 2017), their effects on biotic interactions and community structure and functioning are particularly hard to predict, thus presenting a major challenge for plant ecologists. For priority effects to be useful in invasive species management, further research is needed. First, a better understanding of the mechanisms behind the creation of such priority effects is essential for improving the predictive power of ecology. For instance, although niche preemption by early-arriving species played a role in our study, we cannot exclude the possibility that other niche modification mechanisms, such as the production of allelochemicals by early-arriving species or plant-soil feedbacks, have occurred. Second, because environmental severity affects the strength of facilitative interactions (Brooker *et al.* 2007) and priority effects (Vannette & Fukami 2014; Young *et al.* 2017), additional experiments are needed to determine how the timing of arrival of *S. inaequidens* in native grassland communities and facilitative interactions with natives (either direct or indirect) affect invasion across an environmental stress gradient (e.g., disturbance, resource availability). Finally, priority effects being contingent on environmental conditions during plant establishment (Young *et al.* 2017), we argue that long-term experiments are needed to elucidate how weather conditions during plant establishment affect the strength, direction, and persistence of priority effects (Temperton *et al.* 2016).

## Conclusion

In this study, we showed that the establishment success of an exotic species (*S. inaequidens*) is strongly dependent on its timing of arrival in a European grassland community. Whatever the origin of the species arriving first in the community, they created inhibitory priority effects for species arriving later. *S. inaequidens* benefited more from arriving early than the natives, particularly when it was followed by a mixture of native grasses and legumes. On the other hand, native grassland species created strong inhibitory priority effects that severely affected the growth and N uptake of *S. inaequidens*. Despite some evidence of N facilitation via N sparing in communities containing legumes, we did not find any evidence to support that the presence of this functional group in the community being invaded would favour invasion by *S. inaequidens*. When natives arrived three weeks after *S. inaequidens*, however, the strength of priority effects was lower when legumes were present in the native community. Altogether, our results show that priority effects created by natives probably make the invasion of existing grassland communities by *S. inaequidens* unlikely and are an important mechanism to explain why this exotic species is not commonly found in European grassland communities yet. On the other hand, our results also showed that an early arrival of this exotic species at a site with low native species abundance (e.g., following a disturbance) is a scenario that could potentially favour invasion by *S. inaequidens*. This has direct consequences for conservation and ecological restoration, indicating the need to at all costs avoid creating open space and niches for this species to arrive early and thus establish particularly well into native grassland communities.

## Acknowledgements

We would like to thank Dr Thomas Niemeyer (Leuphana University Lüneburg, Germany) for his invaluable help and technical support. Many thanks also to Carolina Levicek for all the time spent on making the plant illustrations used in this paper (you can have a look at her work here: http://carolinalevicek.com/). This research was funded by the Chair of Ecosystem Functioning and Services, Leuphana University Lüneburg, Germany.

## Author contributions

B.D., E.W. and V.T. conceived the ideas and designed methodology; B.D., E.W. and M.K. performed the experiment; B.D., M.K. and J.N. collected the data; B.D. analysed the data and led the writing of the manuscript. All authors contributed critically to the drafts and gave final approval for publication.

## Data accessibility

Raw data and R scripts used for data analysis are fully accessible here: https://doi.org/10.5281/zenodo.2558204

## Supplementary information

**Table S1.**
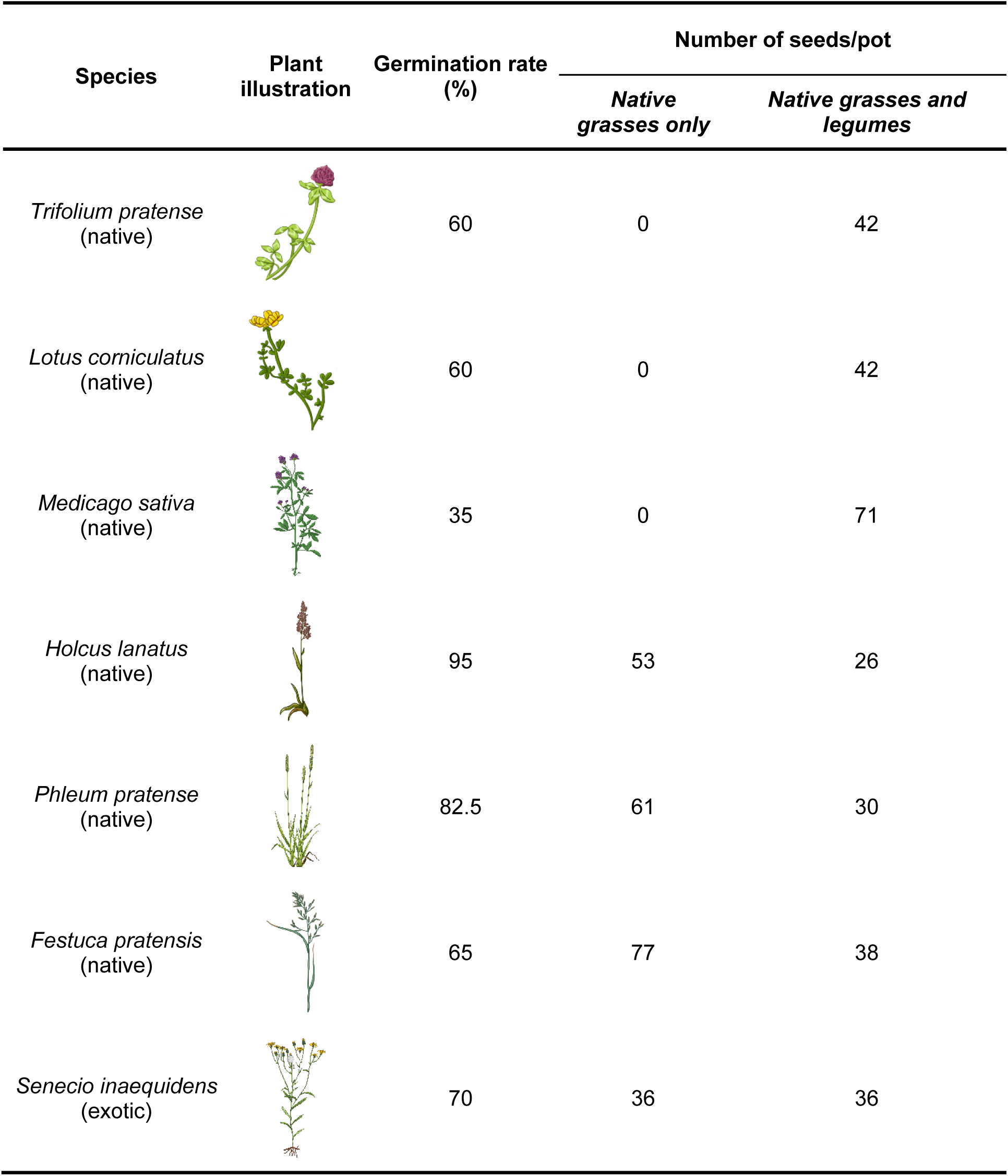
Species composition and sowing densities used in our experiment for each plant community. Plant illustrations by Carolina Levicek.

| Species | Plant illustration | Germination rate (%) | Number of seeds/pot |  |
| --- | --- | --- | --- | --- |
|  |  |  | *Native grasses only* | *Native grasses and legumes* |
| *Trifolium pratense* (native) |  | 60 | 0 | 42 |
| *Lotus corniculatus* (native) |  | 60 | 0 | 42 |
| *Medicago sativa* (native) |  | 35 | 0 | 71 |
| *Holcus lanatus* (native) |  | 95 | 53 | 26 |
| *Phleum pratense* (native) |  | 82.5 | 61 | 30 |
| *Festuca pratensis* (native) |  | 65 | 77 | 38 |
| *Senecio inaequidens* (exotic) |  | 70 | 36 | 36 |

**Figure S1.**
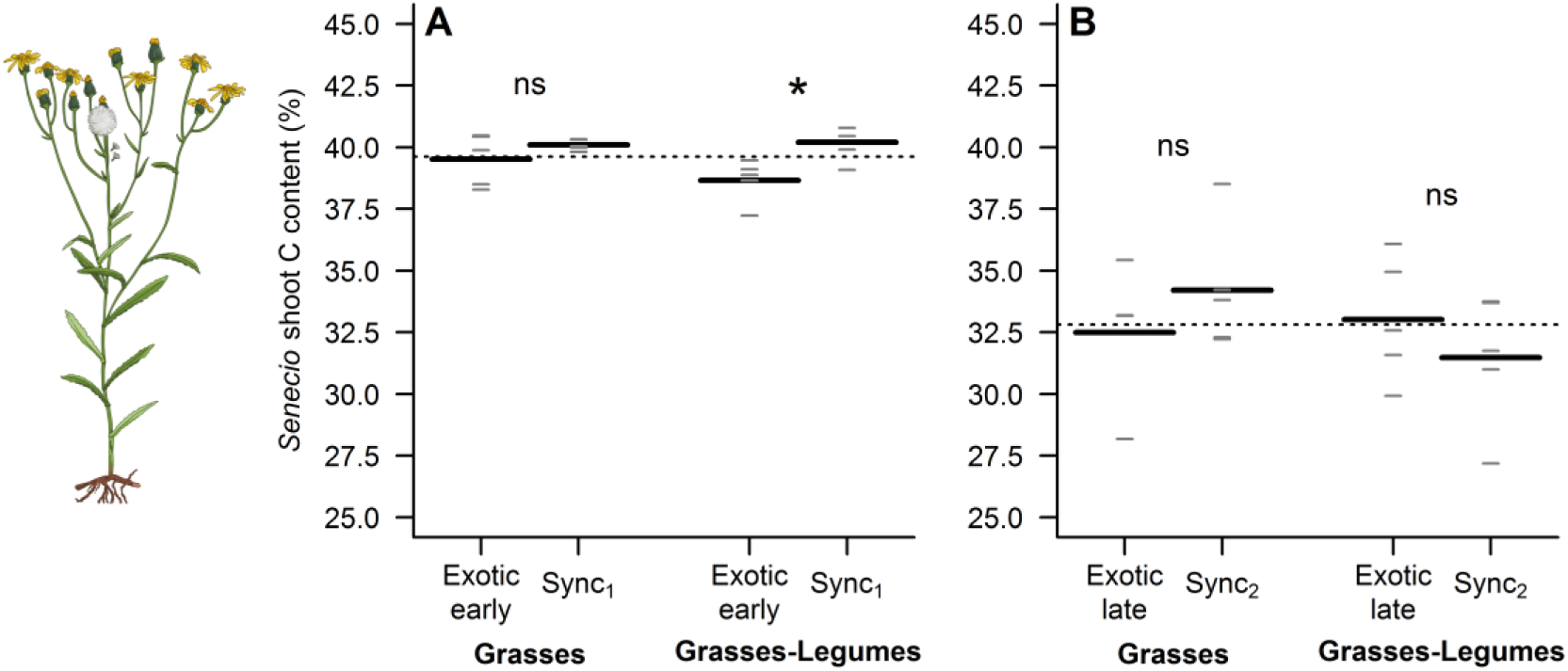
Influence of the timing of arrival of the exotic species and the native plant community composition on the carbon content of *S. inaequidens*. Each graph shows the overall mean (horizontal dotted line), the mean of each group (horizontal black lines, n=4-5), and each individual observation (short horizontal grey lines). Please note that the terms “late” or “early” in all graph always refer to the timing of arrival of *S. inaequidens* (Fig. 1). Because of the low shoot dry weight of *S. inaequidens* obtained for one replicate of the treatment where the exotic species arrived late in a community made of native grass species only, we were not able to measure the C content of that replicate. ns, not significant (P>0.05); *, P<0.05; **, P<0.01; ***, P<0.001.

**Figure S2.**
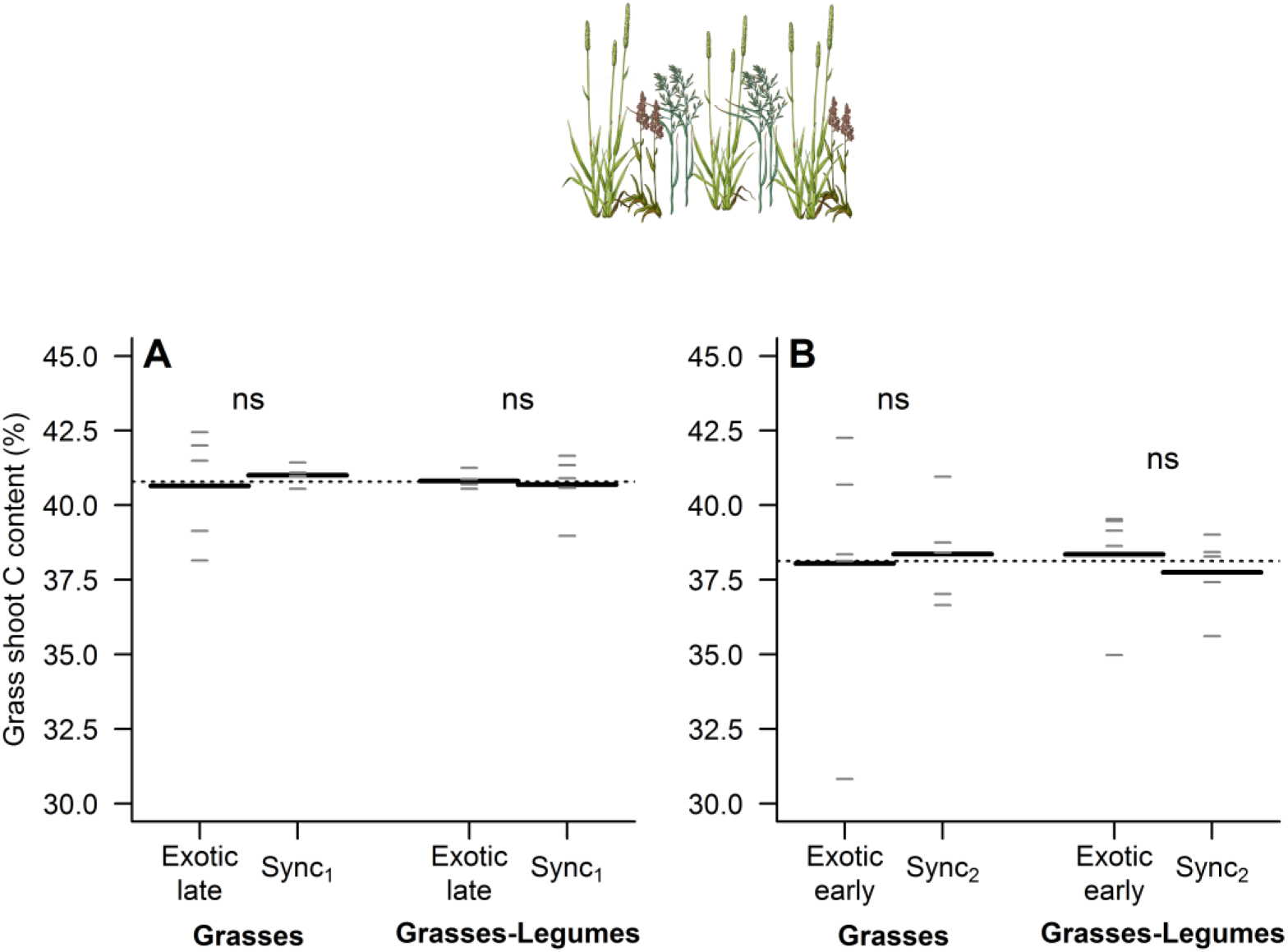
Influence of the timing of arrival of the exotic species and the native plant community composition on the carbon content of the grasses. Each graph shows the overall mean (horizontal dotted line), the mean of each group (horizontal black lines, n=5), and each individual observation (short horizontal grey lines). Please note that the terms “late” or “early” in all graphs always refer to the timing of arrival of *S. inaequidens* (Fig. 1). ns, not significant (P>0.05); *, P<0.05; **, P<0.01; ***, P<0.001.

**Figure S3.**
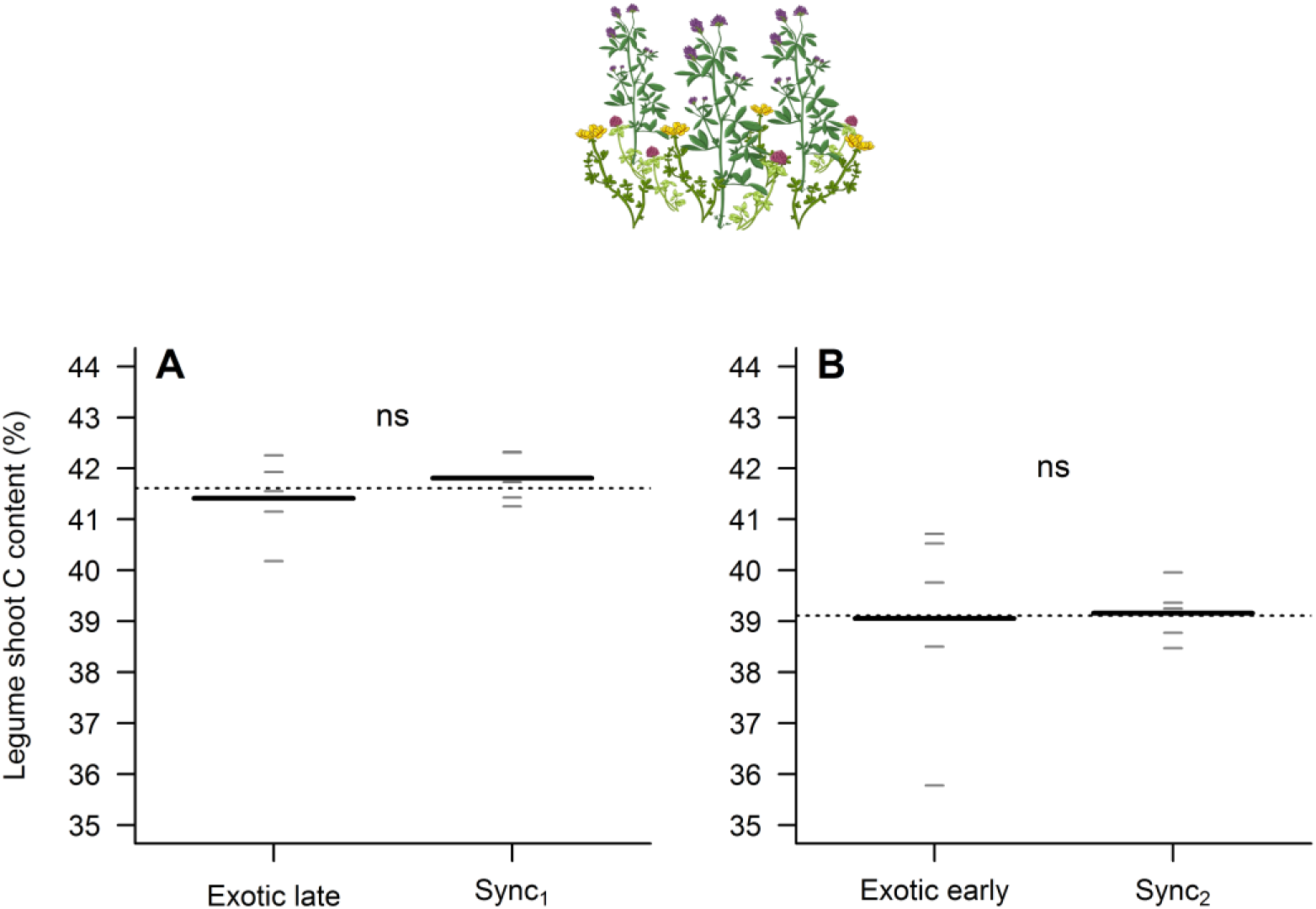
Influence of the timing of arrival of the exotic species on the carbon content of the legumes. Each graph shows the overall mean (horizontal dotted line), the mean of each group (horizontal black lines, n=5), and each individual observation (short horizontal grey lines). Please note that the terms “late” or “early” in all graphs always refer to the timing of arrival of *S. inaequidens* (Fig. 1). ns, not significant (P>0.05); *, P<0.05; **, P<0.01; ***, P<0.001.

**Figure S4.**
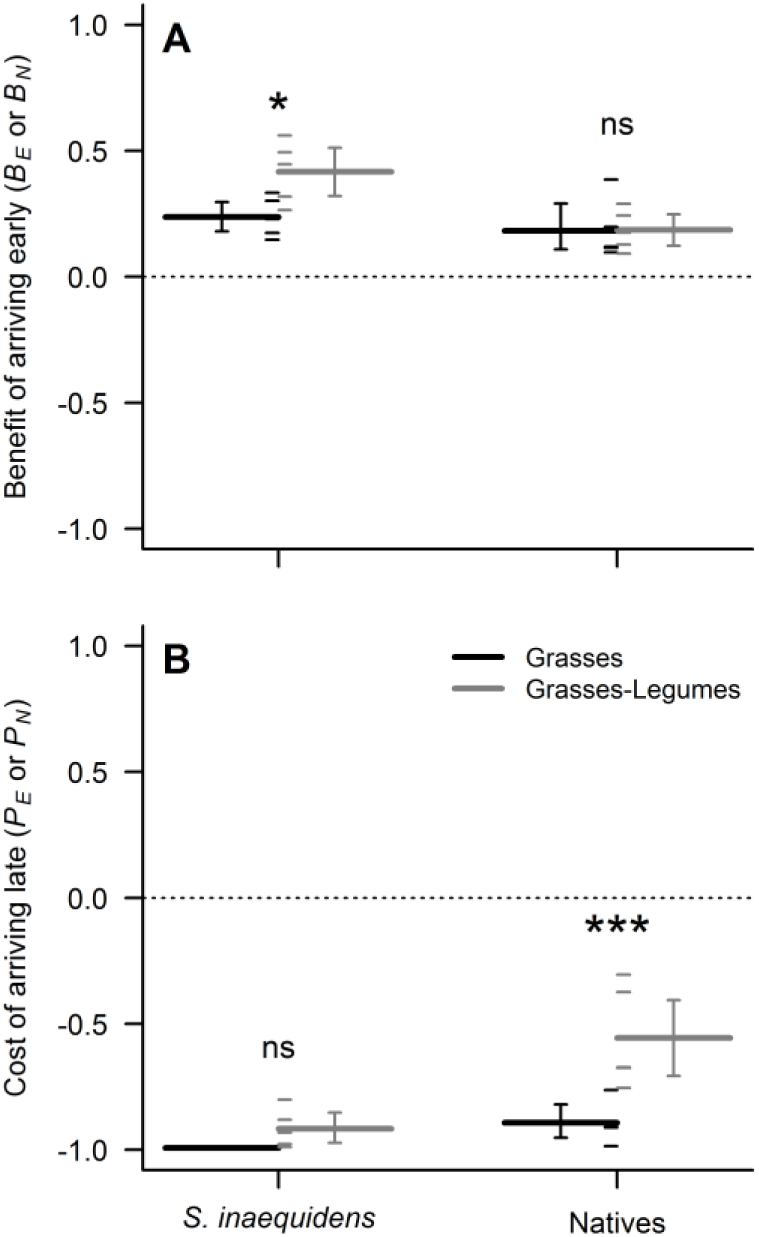
Benefit of arriving early and cost of arriving late in the community for exotic and native species. Benefits and costs were calculated based on shoot N content data using the equations listed in Table 1. All panels show the mean of each group (long horizontal lines, n=4-5), and each individual observation (short horizontal lines). Because of the low shoot dry weight of *S. inaequidens* obtained for one replicate of the treatment where the exotic species arrived late in a community made of native grass species only, we were not able to measure the N content of that replicate. Results are shown separately for each plant species origin (exotic *S. inaequidens* or natives) and each native community composition (see legend). Error bars are 95% confidence intervals computed by bootstrapping using the percentile method. If a 95% confidence interval does not include zero, the mean value of the group is significantly different from zero. ns, not significant (P>0.05); *, P<0.05; **, P<0.01; ***, P<0.001.

**Figure S5.**
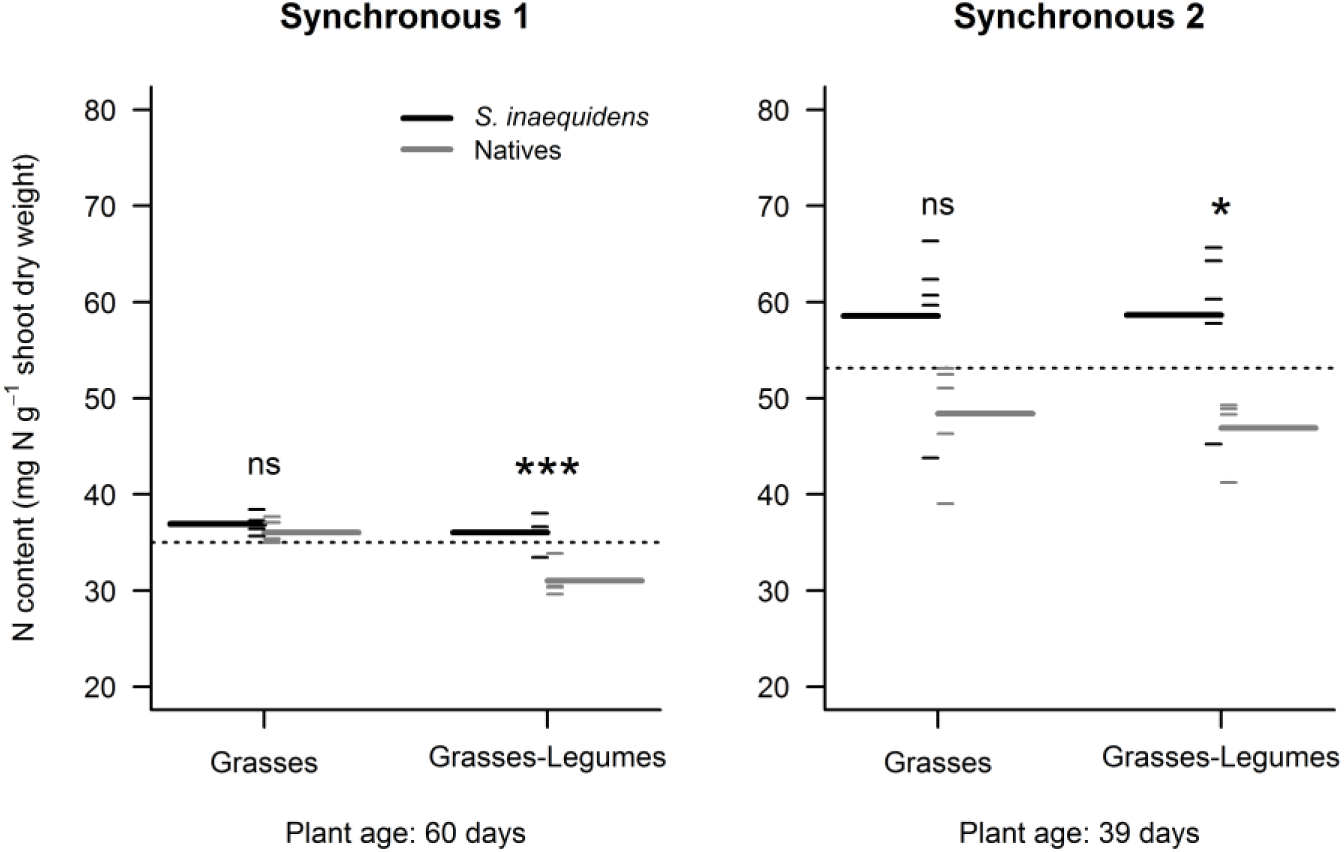
Shoot N content of exotic and native species in plant communities in which all species arrived at the same time. All panels show the overall mean (horizontal dotted line), the mean of each group (long horizontal lines, n=4-5), and each individual observation (short horizontal lines). Results are shown separately for each native community composition (horizontal axis) and each species origin (exotic *S. inaequidens* or natives, see legend). We used the shoot N concentration in the exotic species and the native communities as a proxy for their ability to take up nitrogen from the environment. The results presented here show that *S. inaequidens* took up as much N per unit biomass as a community composed of three native grasses only. When the native community was a mixture of grasses and legumes, however, the N concentration in shoot tissues was on average lower than that measured in the exotic species, thus suggesting a lower ability of the grass-legume mixture to take up nitrogen. The same trends were observed in Synchronous 1 and Synchronous 2 communities. ns, not significant (P>0.05); *, P<0.05; **, P<0.01; ***, P<0.001.

**Figure S6.**
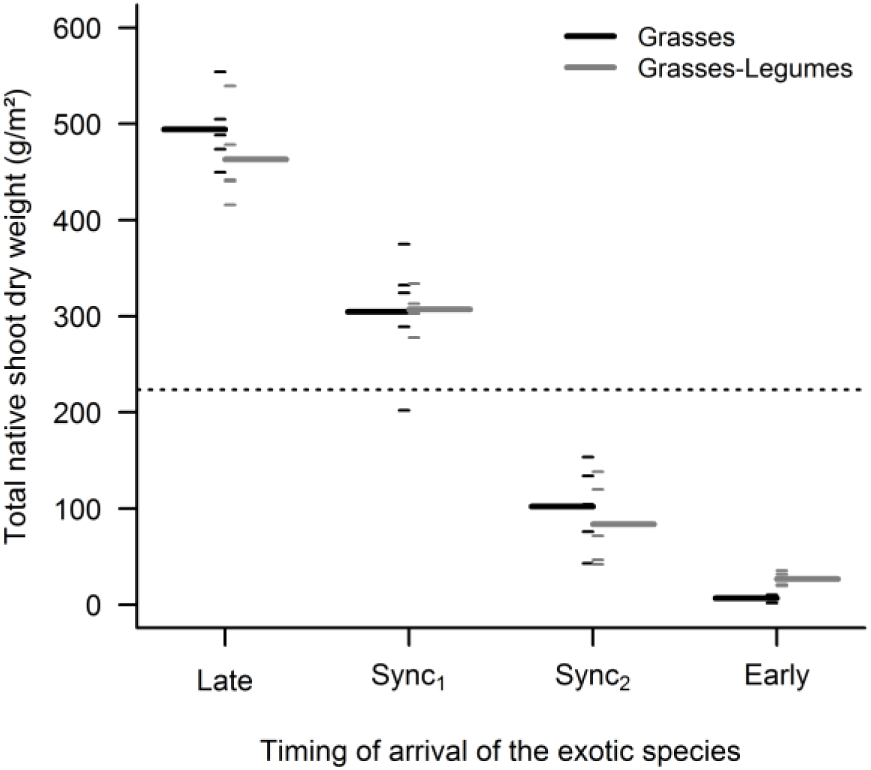
The productivity of the native plant communities did not depend on their species composition. The figure shows the overall mean (horizontal dotted line), the mean of each group (long horizontal lines, n=5), and each individual observation (short horizontal lines). Results are shown separately for each priority effect treatment (horizontal axis) and each native community composition (see legend).

